# From TD_50_ to Benchmark Dose in Nitrosamine Risk Assessment: Evidence from N-Nitrosotrimetazidine Carcinogenicity and TGR Mutation Data

**DOI:** 10.64898/2026.08.26.742063

**Authors:** Mélanie Leheup Foucaud, George Johnson, David Kirkland, Alice Griffon, Richard J. Weaver, Flavia Pasello dos Santos, Stefan O. Mueller

## Abstract

The presence of N-nitrosamine drug substance-related impurities (NDSRIs) in pharmaceuticals represents a significant regulatory and safety challenge due to their classification as “cohort of concern” compounds. This paper describes the toxicological evaluation of N-Nitrosotrimetazidine (NTMZ), performed to refine the initial default acceptable intake (AI) limits of 18–26.5 ng/day established by regulatory authorities.

The evaluation followed a tiered approach: NTMZ was first confirmed as mutagenic *in vitro* via the standard Ames test. To further investigate its genotoxic potential, two *in vivo* studies were conducted in Wistar and transgenic rats. Detection of DNA strand breaks in the liver and duodenum (comet assay) together with positive results in the c*II* mutation assay confirmed an *in vivo* mutagenic mode of action. Benchmark Dose (BMD) analysis of the transgenic rat data yielded a BMDL_50_ of 7 mg/kg/day in the male liver.

To characterize long-term carcinogenic risk, a GLP-compliant 2-year carcinogenicity study was conducted in Wistar rats. Chronic exposure induced dose-dependent increases in liver tumors (hemangiosarcomas, hepatocellular carcinomas and adenomas) and intestinal tumors (adenomas and adenocarcinomas), leading to a Tumor Dose 50 (TD_50_) of 23 mg/kg/day in male rats. Benchmark dose analysis of tumor incidence identified a lowest BMDL_10_ of 2.6 mg/kg/day in females, which served as the basis for deriving an AI of 13 µg/day.

This assessment demonstrates a strong predictive correlation between the BMD derived from the in vivo transgenic model, the BMDL_10_ and the final TD_50_ values obtained in the 2-year carcinogenicity study. These findings provided the scientific basis for establishing a conservative AI of 13 µg/person/day based on the BMDL_10_ and further support the regulatory acceptance and use of BMD-derived approaches for the evaluation of nitrosamine impurities.

## Introduction

N-nitroso compounds are a class of compounds defined by the presence of a nitroso functional group directly attached to a nitrogen atom. This class encompasses N-nitrosamines and other derivatives, such as N-nitrosamides, N-nitrosamidines, N-nitrosoureas, N-nitrosoguanidines, and N-nitrosocarbamates (Loeppky and Michejda, 1994). Recognized as potent mutagenic carcinogens, when present as impurities in drug substances, they fall within the “cohort of concern” as defined in ICH M7(R2) (ICH, 2023). This classification implies that the standard Threshold of Toxicological Concern (TTC) is not applicable and is superseded by the requirement to establish compound-specific Acceptable Intake (AI) limits (Ponting et al. 2024).

In the absence of specific long-term data, regulatory agencies such as the EMA and FDA initially imposed highly conservative default limits (18–26.5 ng/day) for N-nitroso impurities. This framework was later refined using structure-based approaches such as the Carcinogenic Potency Categorization Approach (CPCA), which assigned NTMZ to Category 3, corresponding to an acceptable intake limit of 400 ng/day (EMA, 2024a). However, as these limits are based on structural assumptions rather than empirical biological data, they may not accurately reflect the true carcinogenic potency of Nitrosamine Drug Substance-Related Impurities (NDSRIs). Consequently, the generation of empirical dose-response data was necessary to move beyond theoretical classifications and establish a definitive, compound-specific AI based on demonstrated potency.

NTMZ, identified as an impurity in the piperazine-derived Active Pharmaceutical Ingredient (API) Trimetazidine, exemplifies this need. For chronic therapies, refinement of the AI that moves beyond default classifications is necessary to ensure that acceptable intake limits reflect the compound’s actual biological potency and carcinogenic risk.

The central objective of the work summarized here was to generate a comprehensive toxicological dataset to refine the AI for NTMZ, employing an iterative approach where each phase was designed to address key uncertainties regarding its carcinogenic potential and potency By systematically progressing from *in vitro* assays to *in vivo* mutagenicity studies (including a transgenic rat mutation assay) and concluding with a definitive 2-year GLP-compliant carcinogenicity bioassay, we aimed to fulfill regulatory expectations with the highest level of evidence. This comprehensive approach allowed for the refinement of the initial conservative thresholds into a scientifically justified AI based on demonstrated biological potency.

This tiered analysis is of significant toxicological importance, as it provides a unique opportunity to directly compare Benchmark Dose (BMD) values from short-term mutagenicity models with the TD_50_ and BMDL_10_ derived from chronic exposure in the carcinogenicity bioassay. By supporting the correlation between mutagenic potency (BMD) and carcinogenic outcome (TD_50_ and BMDL_10_), this mutation derived AI approach may contribute to future nitrosamine safety evaluation frameworks. In alignment with 3R principles (Replace, Reduce Refine), our findings underscore the potential for short-term, high-confidence assays to serve as a conservative surrogate for lifetime bioassays, promoting a more ethical and streamlined regulatory process without compromising patient safety.

### Genotoxicity assessment

#### In silico: DEREK and Leadscope

The mutagenicity of NTMZ (Fig.1) was evaluated using a complementary *in silico* approach using two (Quantitative) Structure-Activity Relationship ((Q)SAR) methodologies, in alignment with ICH M7(R2) guideline (ICH, 2023).

**Fig. 1:**
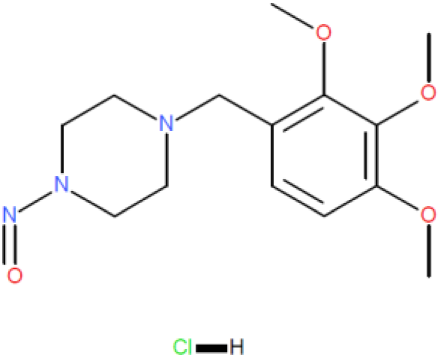
Structure of N-Nitrosotrimetazidine (NTMZ)

An expert rule-based assessment was performed using Derek Nexus (Lhasa Limited). This was complemented by a statistical-based evaluation using Leadscope Model Applier (Instem). Derek Nexus (KB 2025 1.0) concluded that *in vitro* mutagenicity is ’Plausible’. This finding was triggered by the detection of a structural alert for a secondary N-nitrosamine or N-nitramine (Alert 007) which was expected. Using Leadscope (Bacterial Mutation v2), a ’Positive’ prediction for mutagenicity with a high probability of 0.944 was determined. The alignment between these two orthogonal methodologies—expert rule-based and statistical-based— provides a consistent and robust indication of the impurity’s mutagenic potential. In addition, the impurity was assigned to CPCA (Carcinogenic Potency Categorization Approach) Category 3 using Derek Nexus, corresponding to an Acceptable Intake (AI) of 400 ng/day. The CPCA has been adopted by several international regulatory authorities. It assigns N-nitrosamines to potency categories by considering key structural features mapped into an α-hydrogen score, deactivating feature score and activating feature score. A potency score is calculated by summing the contributions from these structural features, according to the following equation

Potency Score = α-Hydrogen Score + Deactivating Feature Scores + Activating Feature Scores

For NTMZ, the resulting potency score of 3 was based on an α-hydrogen score of 1, reflecting the presence of two α-hydrogen atoms on each α-carbon adjacent to the nitrosamine moiety, a deactivating feature score of 2 attributed to the incorporation of the N-nitroso group within a six-membered ring, and an activating feature score of 0.

#### In vitro mutagenicity assay: Ames test

The bacterial reverse mutation test, commonly referred to as the Ames test, is the most widely used assay for evaluating chemically induced gene mutations. Developed by Bruce Ames in the early 1970s (Ames 1971, 1975), this *in vitro* assay utilizes various strains of Salmonella typhimurium and Escherichia coli, specifically designed to detect a broad spectrum of DNA-reactive chemicals, leading to gene mutations.

The Ames test is commonly used as a hazard screening method to evaluate the mutagenic potential of chemicals, pharmaceuticals (EMA 2012) and impurities in pharmaceutical products (ICH, 2023) and acts as a critical predictive surrogate for determining potential *in vivo* genotoxicity and carcinogenicity potential.

##### Study Design and Methodology

To evaluate the mutagenic potential of NTMZ, a standard bacterial reverse mutation assay was conducted in 2019, prior to the widespread implementation of the “Enhanced Ames Test” conditions for nitrosamines (EMA, 2024b). The study was performed in accordance with OECD Test Guideline 471 (OECD 1997).

Five *Salmonella typhimurium* histidine-requiring strains (TA98, TA100, TA1535, TA1537, and TA102) were utilized. NTMZ (CAS No. 92432-50-3, batch number SB-677, purity >99%) and a structural analog, 1,4-dinitrosopiperazine (DNPZ, CAS No. 140-79-4, batch number PH009834) used as a positive nitrosamine control, were dissolved in DMSO. Metabolic activation (S9 mix, 10% v/v) was derived from male Sprague-Dawley rat livers induced by either Aroclor-1254 or Phenobarbital/β-naphthoflavone, the latter being particularly relevant for nitrosopiperazine derivatives (Rao 1978).

Two experimental methods were employed:

Plate Incorporation: tested at seven concentrations ranging from 5 to 5000 µg/plate (triplicate). Pre-incubation: specifically performed for DNPZ (30 min at 37°C) at concentrations from 160 to 5000 µg/plate to enhance the detection of nitrosamine mutagenicity. Treatments of DNPZ were reduced to 0.05 mL due to potential interference as some organic vehicles are known to be near to toxic levels when added at volumes of 0.1 mL in this assay system when employing the pre-incubation methodology.

Three replicate plates for each test concentration were assessed.

Two independent experiments were performed, one using Aroclor 1254-induced S9 and the other using Phenobarbital/β-naphthoflavone-induced S9.

A positive result was defined as a dose-related increase in revertant counts reaching a threshold of ≥1.5-fold (TA102), ≥2-fold (TA98, TA100), or ≥3-fold (TA1535, TA1537) over the concurrent vehicle control.

##### Results

The assay was considered valid as all negative control counts fell within the laboratory’s historical range and positive controls induced the expected significant increases in revertants. NTMZ demonstrated clear mutagenic activity only in the presence of metabolic activation. In strain TA1535 (sensitive for base-pair substitutions), ≥3-fold increases in revertant counts were observed at 500 µg/plate and above with both Aroclor and Phenobarbital/β-naphthoflavone S9. In strain TA100, ≥2-fold increases were observed at 1600 µg/plate and above with Phenobarbital/β-naphthoflavone S9. No mutagenicity was observed in the absence of S9 or in other strains (TA98, TA1537, TA102), although cytotoxicity (reduced background lawn) was noted in TA102 at 5000 µg/plate.

DNPZ (Positive Nitrosamine Control) exhibited a similar mutagenic profile to NTMZ. In the plate incorporation method, increases ≥3-fold were observed in TA1535 at 1600 µg/plate and above with both types of S9. The pre-incubation method significantly increased the assay sensitivity, with positive responses at 625 µg/plate and above in TA1535 and at 5000 µg/plate in TA100 again with both types of S9.

In conclusion, NTMZ was determined to be mutagenic in the presence of metabolic activation, primarily inducing base-pair substitution mutations.

### In vivo genotoxicity assessment

#### Combined in vivo bone marrow micronucleus (MN) and comet assay in rats

##### Rationale and Study Design

Following the positive *in vitro* results, a combined *in vivo* study was conducted to investigate the potential clastogenic/aneugenic activity (Micronucleus test) and DNA strand breakage (Comet assay) of NTMZ. This integrated approach minimizes animal use in alignment with 3R principles while assessing multiple genotoxic endpoints in target and systemic tissues. The study design complied with OECD Guidelines 489 and 474 (OECD 2016a, b). Male Wistar rats (6-7 weeks old) were administered NTMZ hydrochloride (batch SC 401, 98.4% purity) via oral gavage for three consecutive days (0, 24, and 45 hours). Based on a preliminary dose-range finding study (250 to 2000 mg/kg/day), the doses selected for the main study were 56, 111, and 223 mg/kg/day (expressed as base). Since no gender differences in toxicity were seen in the dose-range finding study, only male rats were dosed in the main test. The high dose was selected as the intended Maximum Tolerated Dose (MTD). Vehicle (0.9% NaCl) and positive controls - Cyclophosphamide (19 mg/kg as a single oral dose at 0 hours) for the micronucleus and Ethyl Methanesulfonate (EMS, 200 mg/kg /day as oral doses at 24 and 45 hours) for the comet assay - were included. Five male rats were used for all groups, except for high dose group where 8 rats were administered in case of unexpected death. Blood was sampled in satellite animals at 0.5, 1 and 3h after the last administration for exposure assessment.

##### Tissue Processing and Analysis

Three to four hours after the last administration, tissues of interest, namely liver and duodenum (Comet assay) and bone marrow (micronucleus test) were collected/isolated from 5 animals per group and examined for genotoxic effects. For the Comet assay, cell suspensions from the liver and duodenum were prepared by mincing (liver) or scraping to remove apoptotic cells and then mincing (duodenum). Cell suspensions were mixed with low melting point agarose, and layered on to pre-coated comet slides in triplicate. Cell lysis and DNA unwinding, electrophoresis (20 minutes for duodenum and 30 minutes for liver, which is standard practice in the testing laboratory), and neutralization were carried out according to standard procedures. Slides were then fixed, dried, stained with SYBR Gold and mounted with coverslips. Slides were coded before scoring using fluorescence microscopy and Perceptive Instruments Comet IV image analysis. DNA damage was quantified as % Tail Intensity from 150 cells per tissue per animal. Slides were also examined for presence of severely damaged cells (hedgehogs).

For the Micronucleus test, bone marrow was flushed from the femurs with serum, cells were smeared onto slides, fixed and stained. Although Giemsa staining was used, strict criteria were applied to differentiate micronuclei (MN) from confounding mast cell granules. The proportions of polychromatic and normochromatic erythrocytes (PCE and NCE) were determined as an indicator of bone marrow toxicity. Slides were coded before analysis and micronuclei were scored from at least 4000 PCE per animal, as recommended by OECD Guideline 474 (OECD 2016b). Statistical analyses were performed using ToxRat Professional v3.2.1, with predefined criteria to classify results in both the micronucleus and comet assays. The compound was considered positive if it induced a statistically significant, dose-related increase exceeding historical control limits, and negative if no significant or dose-related effects were observed and all values remained within historical control ranges.

##### Results

The study was considered valid, with negative controls giving micronucleus frequencies and % Tail Intensity values that fell within historical ranges and positive controls inducing significant increases in their respective endpoints. Clinical signs of lethargy, rough coat, and hunched posture were seen in animals dosed at the highest dose of 223 mg/kg. In the high dose group, 2 animals died after the third administration and were replaced by 2 of the additional 3 animals that had been dosed. These clinical observations and the mortality observed at 223 mg/kg confirmed that the Maximum Tolerated Dose (MTD) was reached.

Bone Marrow Micronucleus: NTMZ did not induce a significant increase in the frequency of MN-PCE at any dose level compared to vehicle control (Table 1). The PCE/NCE ratio remained stable across all groups (0.91 to 1.03), indicating a lack of systemic bone marrow toxicity. The results therefore show that NTMZ is not clastogenic or aneugenic in this test system at doses up to the maximum tolerated.

**Table 1.** Summary of the combined in vivo Comet Micronucleus study results.

| Treatment | Dose<br>(mg/kg/day) | Liver<br>% Tail Intensity | Duodenum<br>% Tail Intensity | MN-PCE<br>(per 4,000) | PCE/NCE Ratio |
| --- | --- | --- | --- | --- | --- |
| Vehicle | 0 | 2.91 ± 0.42 | 7.19 ± 1.02 | 3.6 ± 2.9 | 1.01 ± 0.11 |
| NTMZ | 56 | **5.36 ± 1.39\*** | 6.37 ± 2.27 | 3.4 ± 1.9 | 1.00 ± 0.10 |
| NTMZ | 111 | **7.33 ± 2.19\*** | 6.73 ± 1.25 | 5.0 ± 2.3 | 1.03 ± 0.11 |
| NTMZ | 223 | **12.81 ± 4.40\*** | **10.80 ± 2.30\*** | 5.4 ± 2.4 | 0.91 ± 0.19 |
| Positive | EMS/CP | **85.30 ± 2.08\*** | **43.00 ± 2.55\*** | **42.2 ± 22.7\*** | 0.35 ± 0.15 |

\*p < 0.05 vs. vehicle control. EMS: Ethyl MethaneSulfonate. CP: CycloPhosphamide
Bold values: significantly different from corresponding control group (p<0.05)
For micronucleus assay: at least 4000 polychromatic erythrocytes were evaluated with a maximum deviation of 5%.
The PCE/NCE ratio was determined from at least the first 1000 erythrocytes counted.

Comet Assay (Liver and Duodenum): Conversely, NTMZ induced significant DNA strand breakage in target tissues (Table 1). In the liver, a clear dose-dependent and statistically significant increase in % Tail Intensity was observed at all doses (starting from 5.36% at 56 mg/kg/day vs. 2.91% in controls). In the duodenum, a significant increase was observed at the high dose (10.80% vs. 7.19% in controls). No histopathological alterations were noted in these tissues, confirming that the observed DNA damage was not secondary to cytotoxicity.

In conclusion, while NTMZ lacked clastogenic or aneugenic activity in the bone marrow, it is concluded as genotoxic *in vivo*, inducing DNA damage in both the liver and duodenum.

#### In vivo mutation assay at the c*II* locus in Big Blue® transgenic F344 rats

##### Rationale and Study Design

The transgenic rodent (TGR) gene mutation assay, performed in accordance with OECD Guideline 488 (OECD 2025) represents the gold standard for detecting mutations across diverse tissue types. Formally endorsed by the International Workshops on Genotoxicity Testing (IWGT) (Thybaud et al 2003, Heddle et al. 2000) this test is recognized by regulatory authorities as a definitive *in vivo* assessment of mutagenicity, and specifically for nitrosamines (ICH 2011, 2017, 2023). Furthermore, TGR assays demonstrate good correlation with Ames and comet assay (Lambert et al. 2005, Bercu et al. 2023, Jolly et al. 2026).

Big Blue® transgenic rats possess multiple copies of a chromosomally integrated phage shuttle vector (lambda shuttle vector), which incorporates a c*II* reporter gene into the genome of each cell. Mutations are detected by recovering the transgene (c*II* gene) and analyzing the phenotype of the reporter gene in a bacterial host deficient for the reporter gene (OECD 2025).

##### Materials and Methods

Male and female Fischer 344 Big Blue® transgenic rats (F344-TgN (lambda/*lacI*)) were obtained from Taconic Biosciences (Germantown, NY). Animals (8–11 weeks old) were assigned to groups (n = 6/sex/group) and received daily oral administrations for 28 days of either vehicle (0.9% NaCl) or NTMZ (batch SC 455, 98.3% purity). Dose levels were determined based on a 5-day dose-range-finding (DRF) study in which male and female rats received 50, 100, 150, or 200 mg/kg/day of NTMZ. In the DRF assay, NTMZ was well tolerated up to 150 mg/kg/day in males and 100 mg/kg/day in females. Consequently, the following doses were selected for the TGR study: 19, 38, 75, and 150 mg/kg/day for males, and 13, 25, 50, and 100 mg/kg/day for females. Satellite animals (n = 3/sex/group) were included to confirm systemic exposure after the first and last administrations.

DNA isolated from archived frozen tissues of male Big Blue® rats treated with N-ethyl-N-nitrosourea (ENU; 20 mg/kg/day administered on Days 1, 2, 3, 10, 17, and 24) served as a positive packaging control. All animals were necropsied on Day 31 which is acceptable for assessment of mutations in somatic tissues. Liver and duodenum samples from the first five surviving animals per group were processed for DNA isolation and *cII* mutant analysis. Lambda DNA vectors containing the *cII* gene were packaged into phage using Agilent Transpack Packaging Extract, and at least 125,000 phage from at least 2 packagings per tissue sample per animal were analysed for mutations Statistical analysis was performed using a 1-Way Analysis of Variance (ANOVA). A two-sided Dunnett’s Test was performed for pair-wise comparisons of each treated group to the control group.

##### Results

Clinical signs of toxicity, including piloerection and squinted eyes, were observed in males treated at 75 mg/kg/day and above. At the highest dose (150 mg/kg/day), males also exhibited hunched posture and decreased motor activity. In females, hunched posture was also noted at 13 and 50 mg/kg/day. Additional clinical signs of ruffled fur and squinty eyes were observed in females at 50 and 100 mg/kg/day Statistically significant reductions in body weight gain compared to vehicle controls were recorded in males at 38 mg/kg/day (38.3%) and 150 mg/kg/day (36.9%). A non-significant decrease in body weight gain was also observed in treated females.

The positive control (ENU) produced a significant increase in mutant frequency (MF) in both archived frozen tissues validating the sensitivity of the test system for detecting direct-acting mutagens. NTMZ administration resulted in a significant, dose-related increase in *cII* MF in both the liver and duodenum of male and female rats (Table 2). In the liver, significant increases were observed at 38 mg/kg/day and above in males and at the top dose of 100 mg/kg/day in females. In the duodenum, MFs were significantly elevated from all doses in males and at 50 mg/kg/day and above in females. Notably, the MFs that were significantly increased in the duodenum and liver also exceeded the upper 95% confidence limit of the historical negative control range (Table 2).

**Table 2:**
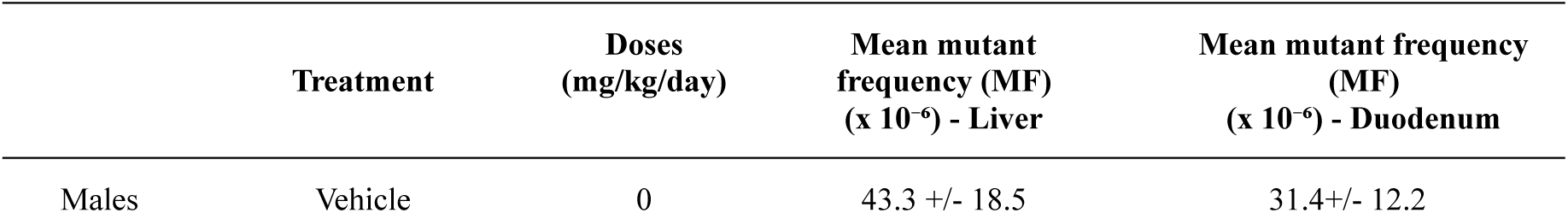
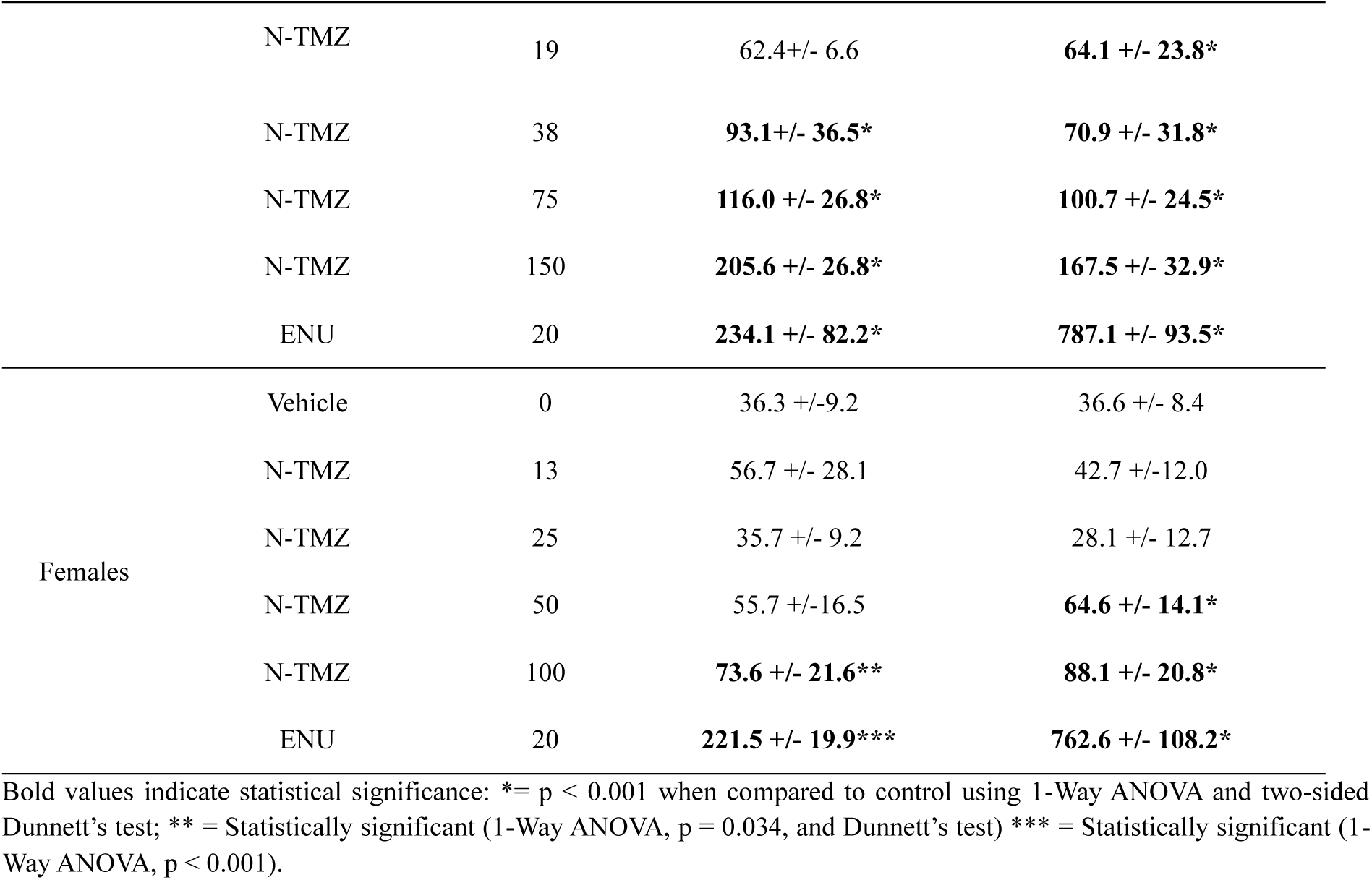
Mean *cII* mutant frequency in liver and duodenum of Big Blue® transgenic F344 rats following 28-day exposure to NTMZ.

The TGR *in vivo* mutation assay results confirm the mutagenic potential of NTMZ previously observed in the Ames and Comet assays. These findings further support the established correlation between these experimental models in genotoxicity assessment (Lambert et al. 2005, Bercu et al. 2023, Jolly et al. 2026, Kirkland 2019).

##### Calculation of Benchmark Dose (BMD)

The Benchmark Dose (BMD) is a statistically derived dose—obtained through mathematical modeling of the full dose-response dataset—that corresponds to a specific level of biological response. It has been widely used for assessing carcinogenicity data (ICH M7(R1) 2017, U.S. EPA 2012, EFSA Scientific Committee 2009) but is increasingly used for assessing genotoxicity data (Jolly et al., 2026, MacGregor et al 2015, Johnson et al., 2014). Apart from the BMD, the BMDL, which is the lower 95% confidence limit of the BMD, is often calculated to serve as a robust Point of Departure (PoD) for risk assessment that accounts for experimental uncertainty (Wheeldon et al. 2021).

To calculate the BMD metrics from the statistical models, a set point on the response-based y-axis must be defined. This is the Benchmark Response (BMR), also referred to as the Critical Effect Size (CES), which carries distinctly different meanings depending on whether it is applied to cancer incidence data or mutation frequency (MF) data. For cancer data, which is inherently quantal and dichotomous, a 10% BMR/CES is typically utilized to estimate an increased incidence of cancer, with a 50% BMR/CES for rodent cancer incidence conceptually analogous to the TD50 (ICH, 2023). In contrast, for mutant frequency data, which is inherently continuous and not quantal, a 50% BMR/CES denotes an increase of 50% above the spontaneous background frequency, an approach more functionally comparable to a No Observed Effect Level (NOEL) for interpretation of mutagenicity data (White et al. 2025).

Covariate BMD analysis was carried out using PROAST v70.0 on the TGR data in liver (Fig.2) and duodenum (Fig.3). The default assumptions in PROAST for covariate BMD analysis were used, with the maximum response (parameter c) and log-steepness (parameter d) assumed equal for all response curves across each study, while parameters for background (a), potency (b) and var (i.e., within group variation) were covariate dependent (Slob and Setzer 2014). Model averaging was used with the default PROAST v70.0 assumptions, with the suite of non-linear models and 200 bootstraps used.

**Fig. 2:**
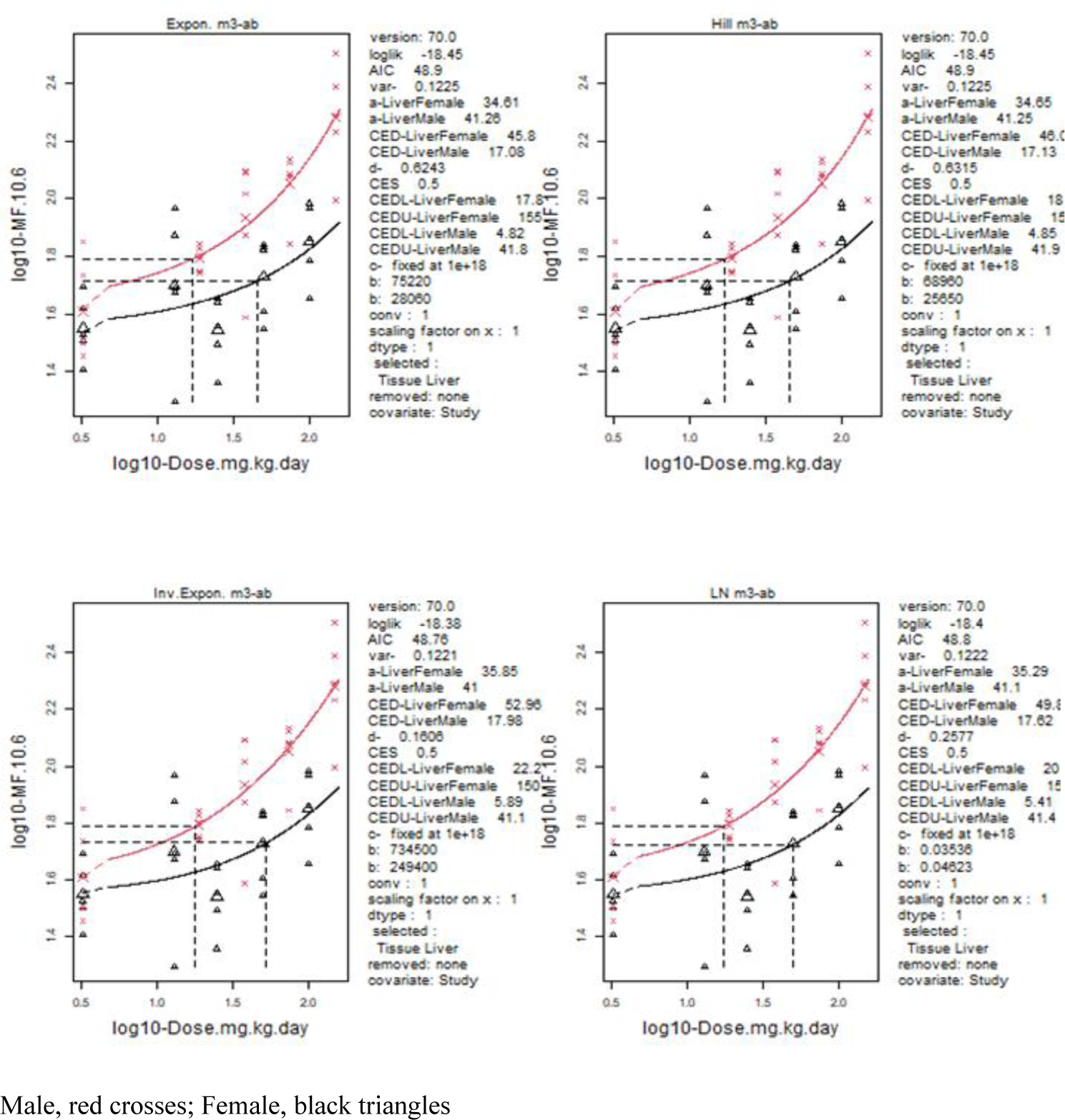
BMD covariate analysis of liver mutation frequency (MF) data for the in vivo TGR mutation assay, using sex as covariate. The 4 standard models used for model averaging in PROAST v70.0 are shown.

**Fig. 3:**
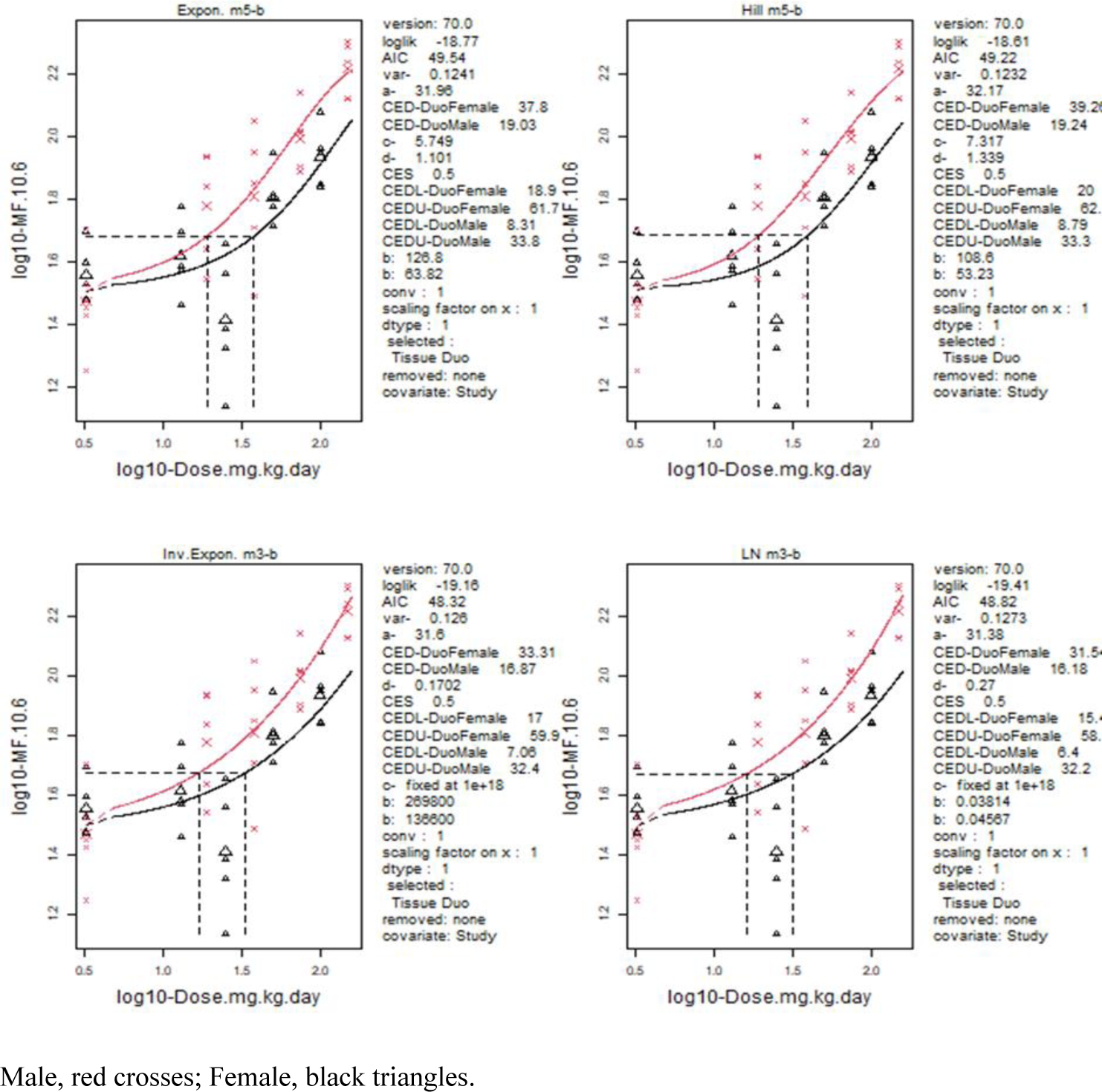
BMD covariate analysis of duodenum Mutant Frequency (MF) data in the in vivo TGR mutation assay, using sex as covariate. The 4 standard models used for model averaging in PROAST v70.0 are shown.

A critical effect size of 50% was used, as recommended by Zeller et al. (Zeller, 2017) and Johnson et al. (Johnson, 2021) and the lower and upper confidence intervals of the BMD (BMDL_50_ and BMDU_50_ respectively) were calculated. The results of the BMD analysis are summarized in Table 3. The BMDL_50_ and BMDU_50_ were calculated for the TGR responses in the liver and duodenum of male and female rats.

**Table 3:** BMD Covariate analysis of the in vivo TGR mutation dose response data from male and female rodents, within duodenum and the liver. Covariate analysis with model averaging was carried out using sex as covariate, with the outputs from the liver analysis in Fig.2 and for duodenum in Fig. 3.

| Covariate BMD | Liver Male<br>(mg/kg/day) | Duodenum Male<br>(mg/kg/day) | Liver Female<br>(mg/kg/day) | Duodenum Female<br>(mg/kg/day) |
| --- | --- | --- | --- | --- |
| BMDL<sub>50</sub> | 7.02 | 7.96 | 14.2 | 19.1 |
| BMDU<sub>50</sub> | 42.8 | 33.8 | 68.1 | 169 |

TGR data were a close fit for a non-linear model, and from Table 3, the most conservative value of the BMDL_50_ was 7.02 mg/kg/day in male liver. The difference between the BMDL_50_ and the BMDU_50_ was always <10-fold which indicates high-quality homogeneous data supporting the robustness of the dose–response dataset

##### Calculating AI from Mutation BMDL_50_

There are ongoing efforts driven by the Genetic Toxicology Technical Committee of the Health and Environmental Sciences Institute (HESI-GTTC), with ongoing discussions with the ICH M7 expert group around the use of mutation data to calculate a cancer-based AI from mutation data. The hypothesis is that mutation potency correlates to cancer potency for nitrosamines, as evidenced by Jolly et al., (Jolly et al. 2026). Using this information, a mutation based BMDL_50_ can be used to calculate an AI through an anchor calculation (establishing a quantitative relationship between a mutagenicity-derived point of departure and a carcinogenicity-derived reference point), or just used within a tabulated series of limits, to provide a more general AI. Using the anchor-based approach, as detailed in Jolly et al (2026) as

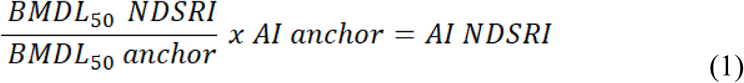

the AI ranges from 1855 to 8802 ng/day (Table 4), and using the category approach of Jolly et al., (2026) which has a pre-set range of BMDL_50_ values that correspond with a pre-defined AI limit, the AI is 1500 ng/day.

**Table 4:** Anchor calculation, to derive AI from in vivo mutation TGR data for NTMZ, by extrapolation from mutation BMDL and AI from other nitrosamines. BMDL for other nitrosamines are taken from Jolly et al., 2026.

|  | Published AI (ng/day) | BMDL<sub>50</sub> (mg/kg/day) | BMDU<sub>50</sub> (mg/kg/day) | Ratio BMDL<sub>50</sub> NTMZ/anchor | Predicted AI of NTMZ by anchor-based approach (ng/day) |
| --- | --- | --- | --- | --- | --- |
| NDMA | 96 | 0.21<sup>a</sup> | 0.32<sup>a</sup> | 33 | 3200 |
| NDEA | 26.5 | 0.03<sup>b</sup> | 0.58<sup>b</sup> | 233 | 6183 |
| NDEA | 26.5 | 0.1<sup>c</sup> | 1<sup>c</sup> | 70 | 1855 |
| NDELA | 1900 | 6.6<sup>a</sup> | 17.3<sup>a</sup> | 1 | 2015 |
| NMOR | 127 | 0.1<sup>b</sup> | 0.71<sup>b</sup> | 69 | 8802 |
| NTMZ | 400 | 7.02<sup>c</sup> | 42.8<sup>c</sup> | 1 | - |

NDMA: N-Nitrosodimethylamine, NDEA: N-Nitrosodiethylamine, NDELA : N-Bis(2-hydroxyethyl) nitrous amide (N-nitrosodiethanolamine, NMOR : N-Nitrosomorpholine, NNK : Nicotine derived nitrosamine ketone or 4- (methylnitrosamino)-1-(3-pyridyl)-1-butanone
<sup>a</sup>: BMD obtained from MutaMouse study; <sup>b</sup>: BMD obtained from BigBlue® Mouse; <sup>c</sup>: BMD obtained from BigBlue® Rat

#### Carcinogenicity study

##### Rationale

Although Benchmark Dose (BMD) modeling is increasingly discussed in scientific literature, it has not yet achieved full regulatory acceptance as the primary basis for risk assessment in this context. According to the latest revisions of the EMA guideline on nitrosamines (EMA 2024), the calculation of a TD_50_ derived from a robust 2-year carcinogenicity bioassay remains the “gold standard” for establishing potency categories and acceptable intake (AI) limits. Consequently, the 2-year study described below was initiated to provide a definitive regulatory basis for the safety assessment of NTMZ, ensuring compliance with current international regulatory expectations for lifetime risk characterization.

##### Materials and Methods

NTMZ was evaluated in a 2-year GLP-compliant carcinogenicity bioassay. Wistar Han rats (n = 50/sex/group) received daily oral gavage administrations of NTMZ (batch SC 606, purity >99%) at 0 (0.9% NaCl control), 2, 6, 15, and 40 mg/kg/day for up to 104 weeks. Doses were selected based on the TGR study results described above: the low doses (2 and 6 mg/kg/day) were anticipated to be non-mutagenic and non-carcinogenic; the high dose (40 mg/kg/day) was expected to be clinically tolerated despite its demonstrated *in vivo* mutagenicity; and the intermediate dose (15 mg/kg/day) was included to help characterize the dose-response relationship for potential neoplastic effects. Animals (8 weeks old at start) were randomly allocated to the various groups and monitored under standard husbandry conditions, with predefined criteria for clinical observations, body weight, food consumption, and survival. Blood samples were collected for toxicokinetics (Weeks 4, 26, 64 and 78) and clinical pathology (Week 51), and systemic exposure was confirmed across all dose levels. At scheduled or unscheduled necropsy, comprehensive macroscopic examination and histopathological evaluation of tissues were performed, with tumour incidence analysed statistically in accordance with ICH/OECD/FDA guidance (ICH S1B, 1995, OECD 451, 2018, FDA, 2001, 2003).

##### Results and conclusion

A treatment-related increase in mortality was observed in both sexes across all dose groups, exhibiting a statistically significant dose-dependent trend. This led to the early termination of treatment of different groups: weeks 67 and 82 for males and females at 40 mg/kg/day, and weeks 96 and 97 for those at 15 mg/kg/day, respectively. Consequently, animals in the 40 mg/kg/day group were sacrificed at week 71 (males) and week 86 (females), while males in the 15 mg/kg/day group were sacrificed at week 101/102. This mortality was associated with a higher occurrence of tumors in the liver (both sexes) and intestines (males only) observed at 15 mg/kg/day and above.

Pathological examination (Table 5) revealed a related increased incidence of hemangiosarcomas in liver in males and females dosed at 6 mg/kg/day and above, and of hepatocellular adenomas and carcinomas at ≥ 15 mg/kg/day in both sexes, correlating with an increased frequency of hepatic masses noted during necropsy. Non-neoplastic proliferative changes, specifically foci of cellular alteration, were identified in the liver across all treated groups.

**Table 5:**
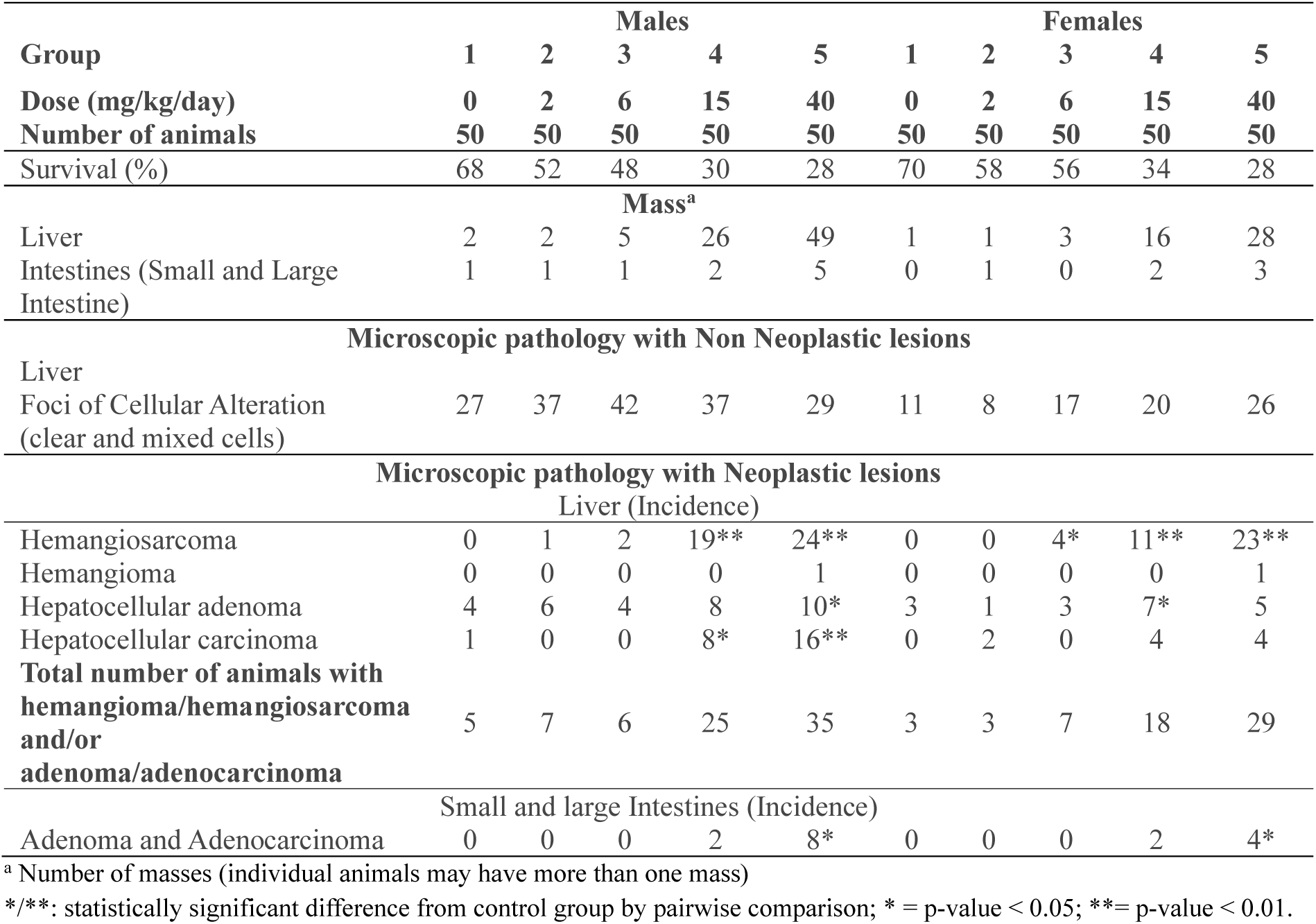
Summary of the main carcinogenicity pathology results.

In the gastrointestinal tract, the incidence of adenomas and adenocarcinomas in both the small and large intestines was significantly elevated in both sexes at doses ≥ 15 mg/kg/day, which also correlated with a slightly higher frequency of macroscopic intestinal masses.

##### TD_50_ and BMD calculation

The TD_50_ (tumorigenic dose 50) is a numerical measure of the carcinogenic potency of a chemical agent. It was formally defined by Peto et al. in 1984 (Peto et al. 1984, Sawyer et al. 1984). In essence, the TD_50_ represents the chronic dose rate (expressed in mg/kg body weight/day) that would halve the probability of an animal remaining tumor-free throughout its standard lifespan, after correcting for mortality from other causes (Gold et al. 1984).

Based on these neoplastic findings, TD_50_ values were calculated to identify the most sensitive site of tumor induction (Table 6), in accordance with ICH M7(R2) guidelines (ICH 2023). Calculations were performed using the Thresher TD₅₀ model (Model 2) implemented within R (https://cran.r-project.org) (Thresher et al. 2020).

**Table 6:**
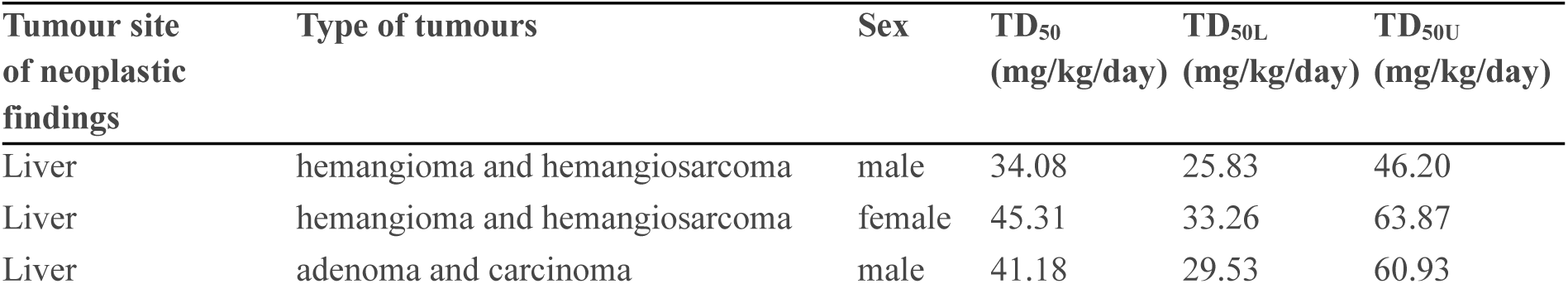
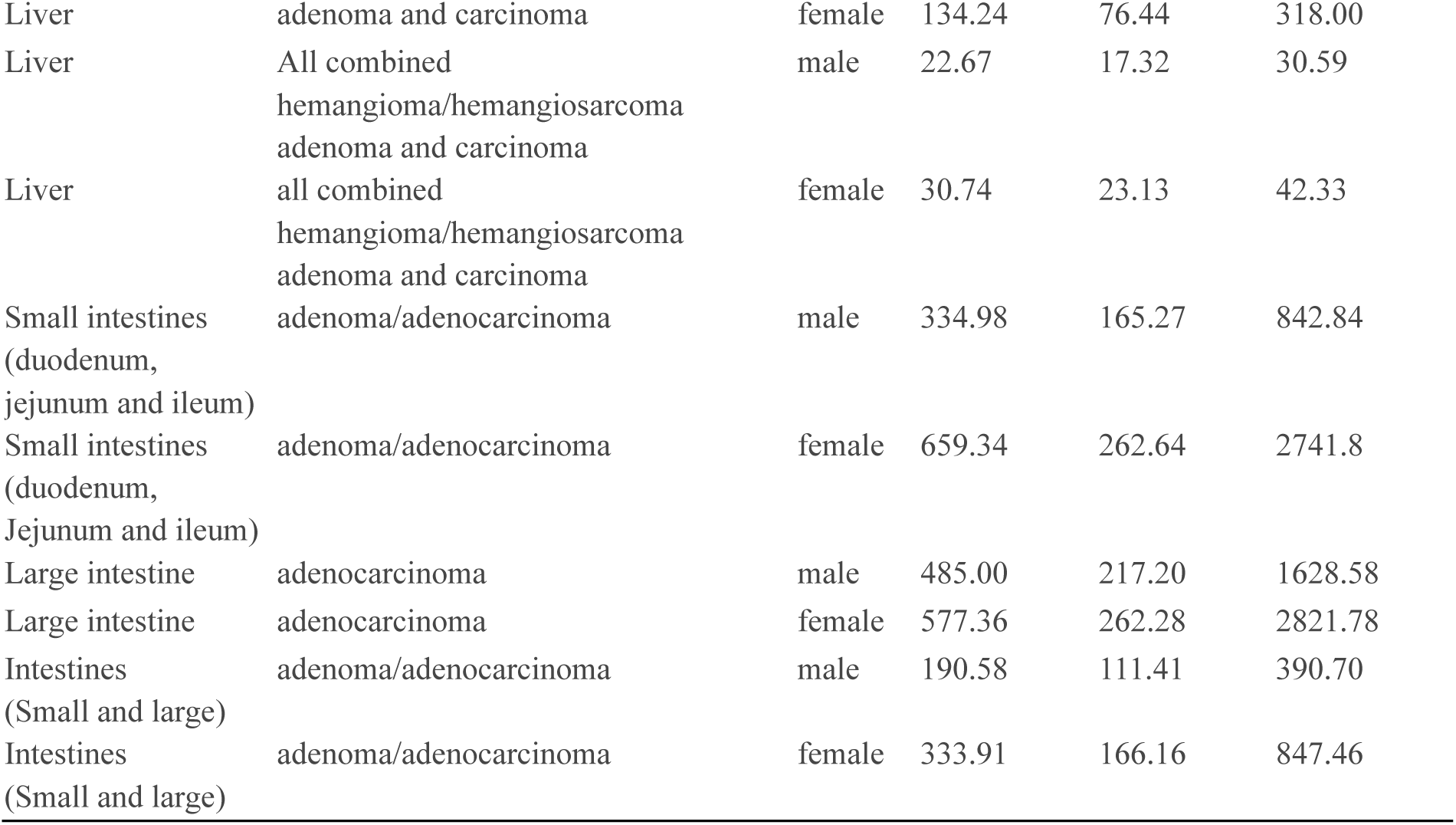
TD_50_ Values for NTMZ.

| Tumour site of neoplastic findings | Type of tumours | Sex | TD<sub>50</sub> (mg/kg/day) | TD<sub>50L</sub> (mg/kg/day) | TD<sub>50U</sub> (mg/kg/day) |
| --- | --- | --- | --- | --- | --- |
| Liver | hemangioma and hemangiosarcoma | male | 34.08 | 25.83 | 46.20 |
| Liver | hemangioma and hemangiosarcoma | female | 45.31 | 33.26 | 63.87 |
| Liver | adenoma and carcinoma | male | 41.18 | 29.53 | 60.93 |
| Liver | adenoma and carcinoma | female | 134.24 | 76.44 | 318.00 |
| Liver | All combined<br>hemangioma/hemangiosarcoma<br>adenoma and carcinoma | male | 22.67 | 17.32 | 30.59 |
| Liver | all combined<br>hemangioma/hemangiosarcoma<br>adenoma and carcinoma | female | 30.74 | 23.13 | 42.33 |
| Small intestines<br>(duodenum,<br>jejunum and ileum) | adenoma/adenocarcinoma | male | 334.98 | 165.27 | 842.84 |
| Small intestines<br>(duodenum,<br>Jejunum and ileum) | adenoma/adenocarcinoma | female | 659.34 | 262.64 | 2741.8 |
| Large intestine | adenocarcinoma | male | 485.00 | 217.20 | 1628.58 |
| Large intestine | adenocarcinoma | female | 577.36 | 262.28 | 2821.78 |
| Intestines<br>(Small and large) | adenoma/adenocarcinoma | male | 190.58 | 111.41 | 390.70 |
| Intestines<br>(Small and large) | adenoma/adenocarcinoma | female | 333.91 | 166.16 | 847.46 |

The liver was identified as the most sensitive and relevant organ, with a TD_50_ of 34 and 45 mg/kg/day for the combined incidence of hemangiomas and hemangiosarcomas in males and females, respectively. When considering the total incidence of hepatic tumors of any origin (vascular: hemangioma/hemangiosarcoma; hepatocellular: adenoma/adenocarcinoma), the calculated TD_50_ was 23 and 31 mg/kg/day for males and females, respectively.

The AI was derived from the TD_50_ according to the following equation:

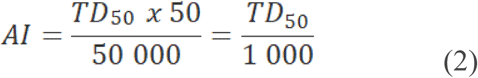

Applying this most conservative TD_50_ of 23 mg/kg/day (obtained from all combined hemangioma, hemangiosarcoma, adenoma and carcinoma) result in a calculated AI of 23 µg/person/day (23 000 ng/day).

BMD analysis (Fig.4) was also carried out on all combined hemangioma, hemangiosarcoma, adenoma and carcinoma data in liver. The results of the BMD analysis are summarised in Table 7.

**Fig. 4:**
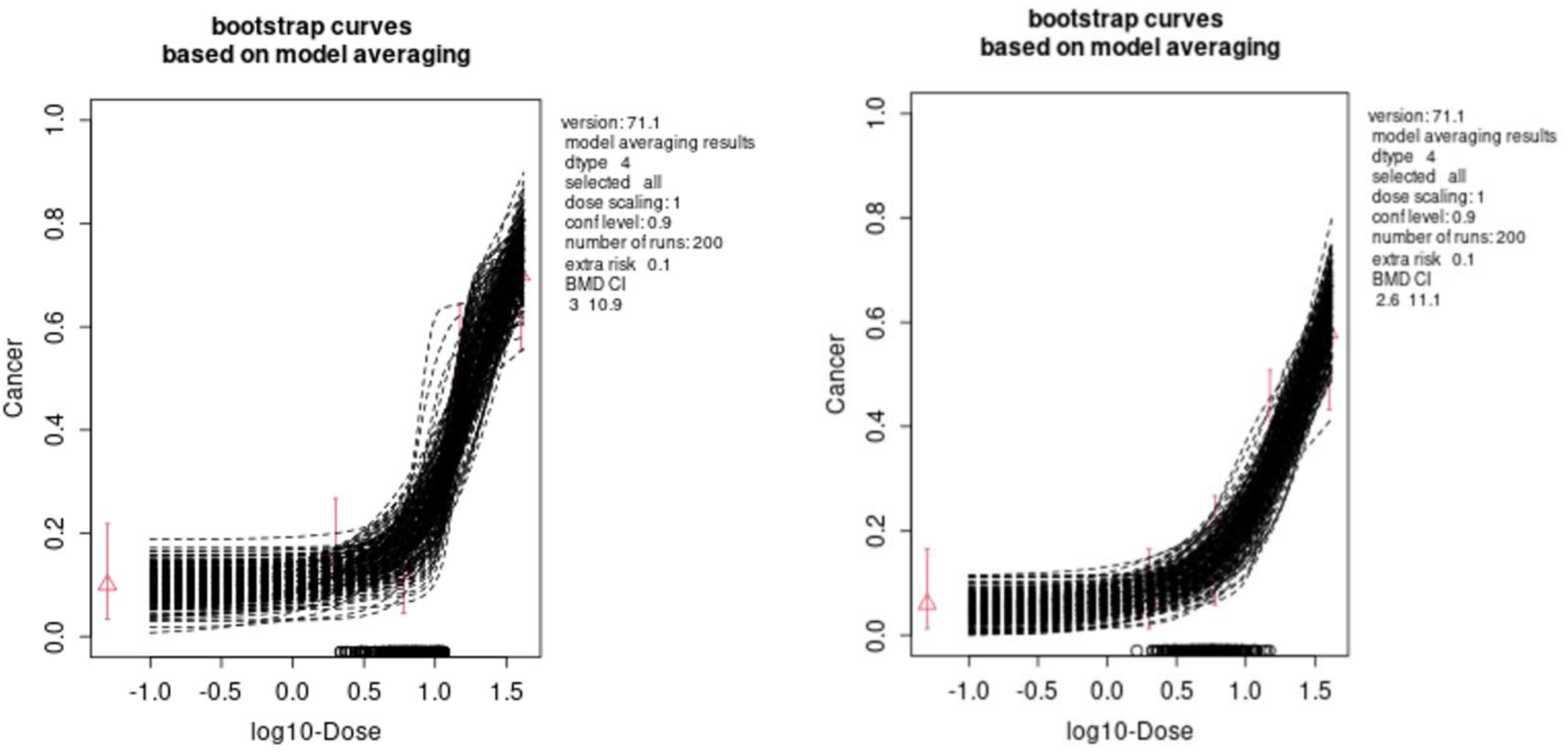
BMD analysis of cancer bioassay data for total incidence of hepatic tumors of any origin in male (left graph) and female (right graph) Wistar rats.

**Table 7:** BMD analysis of the carcinogenicity dose response data from all combined hemangioma, hemangiosarcoma, adenoma and carcinoma of male and female rodents

|  | **BMDL<sub>10</sub>** | **BMDU<sub>10</sub>** | **BMDL<sub>50</sub>** | **BMDU<sub>50</sub>** |
| --- | --- | --- | --- | --- |
| Male | 3 | 11 | 14 | 27 |
| Female | 2.6 | 11 | 21 | 40 |

Applying the most conservative BMDL_50_ of 14 mg/kg/day results in a calculated AI of 14 µg/person/day (14 000 ng/day). The BMD_50_ has a benchmark response (BMR) of 50%, which is comparable to the TD_50_. However, the BMR of 10% can also be used for risk assessment purposes. Therefore, the BMDL_10_ of 2.6 mg/kg/day can be used to calculate the AI at 13 µg/person/day (13 000 ng/day) which represents a more conservative approach compared to the TD_50_-based AI derivation.

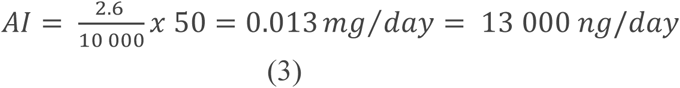

## Discussion

The toxicological evaluation of N-Nitrosotrimetazidine (NTMZ) presented here represents a comprehensive, data-driven strategy to refine safety limits for Nitrosamine Drug Substance-Related Impurities (NDSRIs). By progressing from *in silico* predictions to a definitive 2-year carcinogenicity bioassay, this investigation provides a rare and robust empirical assessment of the tiered approach for assessing compounds within the “cohort of concern” (ICH 2023).

A pivotal outcome of this evaluation is the observed concordance between short-term in vivo mutagenicity endpoints and long-term carcinogenic outcomes. Mutagenic responses were observed *in vivo* in both the Comet assay and the Big Blue® transgenic rodent (TGR) mutation assay, following initial *in vitro* results from the Ames test.

Specifically, the Benchmark Dose (BMD) analysis derived from the TGR study yielded a BMDL_50_ of 7.0 mg/kg/day for the male rat liver. Remarkably, this point of departure from a 28-day study aligns closely with the TD_50_ of 23 mg/kg/day obtained from the 104-week bioassay. The liver and duodenum exhibited treatment-related effects across all *in vivo* models, supporting the hypothesis that site-specific mutagenicity may serve as an early indicator of subsequent neoplasia (hemangiosarcomas and hepatocellular carcinomas). This correlation supports the hypothesis that for nitrosamines, mutagenic potency is a direct driver of carcinogenic potency (Jolly et al. 2026; Kirkland 2019). The initial regulatory landscape for NTMZ was characterized by high uncertainty, with default AI limits ranging from 18 ng/day (EMA) to 400 ng/day (CPCA Category 3). While the CPCA provided a necessary framework for class-based prioritization, our results demonstrate that structural assumptions can significantly overestimate the risks defined by actual biological safety profile of complex nitrosamines (NDSRIs). The empirical TD_50_ of 23 mg/kg/day derived from the 2-year GLP study justifies an AI of 23 µg/day and the conservative BMDL_10_ of 2.6 mg/kg/day justifies an AI of 13 µg/day —nearly 1,300 and 700 times higher than the initial default limit of 18 ng/day, respectively and 57 and 32 times higher than the CPCA limit, respectively.

While the 2-year bioassay remains the “gold standard” (Peto et al. 1984; EMA 2023), it is resource-intensive and consumes a large number of animals. Our data contribute to the growing consensus within the HESI-GTTC and ICH M7 expert groups that high-confidence mutagenicity data (TGR BMDL) can serve as a conservative and scientifically sound surrogate for setting AI limits. In our assessment, the BMDL_50_ from the TGR study (7.0 mg/kg/day) provided a conservative estimate that, if used with appropriate “anchor” calculations (Thresher et al. 2020), would have yielded an AI within the same order of magnitude as that ultimately derived from the carcinogenicity study. This aligns with 3R principles (Replacement, Reduction, Refinement) and suggests a path forward where complex nitrosamines could be regulated based on robust 28-day TGR studies when suitable read-across is unavailable.

The stepwise evaluation strategy integrating *in silico* predictions, *in vitro* mutagenicity data, *in vivo* genotoxicity studies, and long-term carcinogenicity data demonstrated a high degree of concordance across the different levels of assessment, supporting derivation of the final AI of 13 μg/day from the BMDL_10_ (Fig.5).

**Fig. 5:**
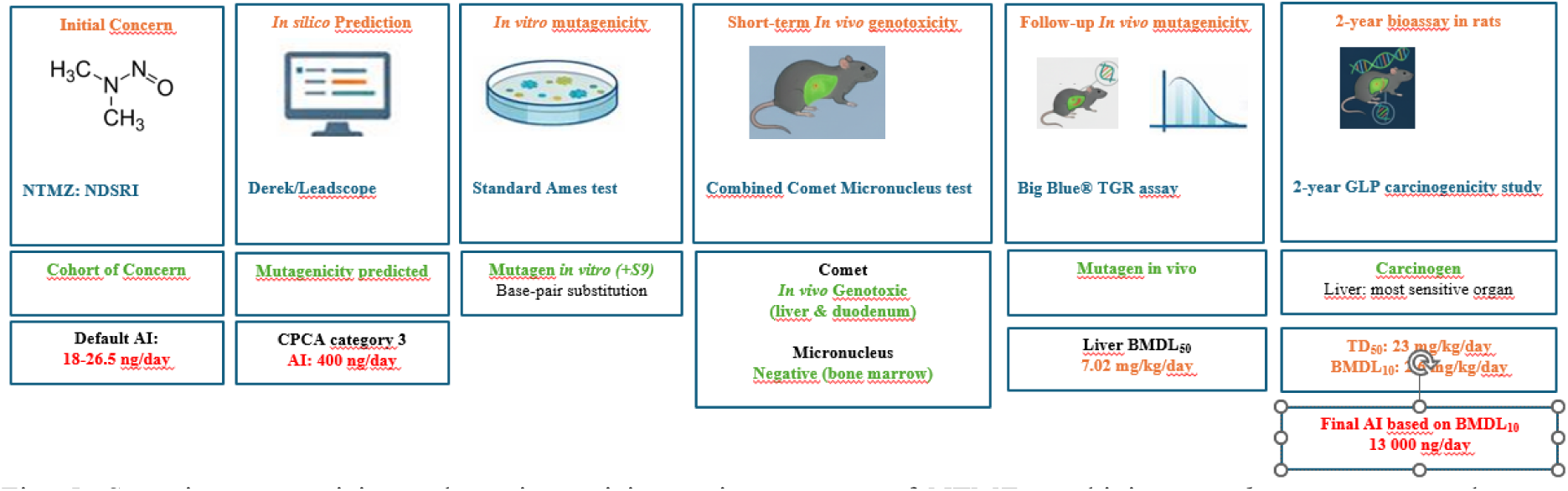
Stepwise genotoxicity and carcinogenicity testing strategy of NTMZ combining *in silico, in vitro,* and *in vivo* approaches.

## Conclusion

N-Nitrosotrimetazidine (NTMZ), a complex N-nitrosamine drug substance-related impurity (NDSRI), was shown to be mutagenic *in vivo* while exhibiting a carcinogenic potency considerably lower than that predicted by conservative default approaches.

The acceptable intake (AI) of 13 µg/day derived from the most conservative carcinogenicity benchmark dose (BMDL_10_) provides a scientifically justified basis for risk assessment and regulatory decision-making.

Importantly, the relationship observed between the TGR-derived BMDL_50_ and the carcinogenicity-derived TD_50_ and BMDL_10_ further supports the growing evidence that *in vivo* mutagenicity data can provide meaningful insight into the carcinogenic potency of NDSRIs. Beyond the assessment of NTMZ itself, these findings contribute to ongoing regulatory discussions on benchmark dose-based approaches and support their further consideration within the evolving ICH M7 framework as a science-based strategy for the evaluation of complex nitrosamines.

## Supplementary Information

## Acknowledgement

We would like to express our deepest gratitude to Françoise Goldfain-Blanc for her unvaluable contribution in defining and guiding the strategy for evaluating the risk of this nitrosamine.

## Author contributions

**Mélanie Leheup Foucaud:** Conceptualization, Data curation, Investigation, Writing -original draft.

**George Johnson**: Investigation, Writing-original draft, review and editing

**David Kirkland**: Writing: review and editing

**Alice Griffon:** Writing: review and editing

**Richard J. Weaver:** Writing: review and editing

**Flavia Pasello dos Santos**: Writing: review and editing

**Stefan O. Mueller:** Funding Acquisition, Writing: review and editing

## Ethical Approval

All animal studies were conducted in accordance with relevant national and international guidelines for the care and use of laboratory animals and were approved by the appropriate institutional ethics committees.

## Informed consent

Not applicable.

## Data Availability

The data supporting the findings of this study are available from the corresponding author upon reasonable request and subject to applicable confidentiality requirements.

